# Memory for a day: Olfactory place episodic-like memory for dung droppings by the domestic horse (*Equus ferus caballus*)

**DOI:** 10.1101/2022.11.01.514774

**Authors:** Audrey EM Guyonnet, Ian Q Whishaw

## Abstract

The domestic horse (*Equus ferus caballus*) makes dung deposits to form “stud-piles” and investigates dung droppings more generally, suggesting that dung contains species-relevant communicative information. This natural behavior provides a behavior with which to examine a species-typical form of memory used by horses in relation to their social behavior. Horses were video recorded in indoor and outdoor riding arenas as they were lightly taken on a lead line to experimenter-determined objects or dung-piles. Frame-by-frame video analysis was used to measure sniffing duration and spatial memory of dung dropping visitations. Horses readily approached and sniffed dung for longer durations than they sniffed other objects. They always approached dung at new locations, made head movements across the extent of dung-piles as they sniffed, showed no preference in the nostril directed to the target, and might blink during sniffing and always blinked when disengaging from sniffing. Horses did remember dung visited withing a day by reducing visits and sniff duration but displayed little retention between days. The results are discussed in relation to the idea that this form of episodic-like memory is time limited because it competes with safety-related behavior related to horse movement within foraging areas.

## Introduction

Olfactory signalling is a widely used communication strategy of vertebrates. In addition to body odor, animals use specialized glandular exudates, signals from saliva, and signals from urine and feces. Signalling may be through the air, object marking, self-marking, social partner marking and in the case of urine and feces, incidental marking. For horses (*Equus ferus caballus*), olfactory signals may reveal information about species, group identification, sex, reproductive status, age, mood and personal identification (Eisenherg and Kleitman, 1972; Hothersall et al, 2010; Marinier et al, 1988). Previous studies report that the horse engages in interindividual communication through both incidental and intentional deposition of urine and dung (Klingel, 1975; Welsh, 1975; Feist and McCullough, 1976; Berger, 1987; Turner et al. 1981; Salter and Hudson, 1982; Rubenstein and Hack, 1992). Taken together, this research suggests that horses can organize their spatial and social communication via the odor of urine and dung (King and Gurnell, 2006; Krueger and Flauger, 2011; Péron et al., 2014). For example, stallions repeatedly defecate in one place to form large stud-piles in the regions that a herd occupies. Subsequently, the depositor and other horses encountering a stud pile may sniff the pile before adding to it, and while adding to it may disturb the pile by walking through it thus distributing its scent (Feist and McCullough, 1976; McDonnell, 2003; Saslow, 2002). Stallions also deposit feces in locations where females urinate, presumably to negate signals that mares might send to a competitor (Linklater, 2000; Salter and Hudson, 1980; Turner et al., 1981). The purpose of scent marking is often discussed as signalling the presence, status and competitive ability of a resident stallion in relation to female resources and by females to signal their location and reproductive status (Rubenstein and Hack, 1992).

Scent marking behavior and olfactory investigation of feces suggests that horses have a good memory for the areas they occupy and that odor-based signalling may be part of their spatial ecology, i.e., they use odor to update their frame of reference with respect to their movement (Feist and McCullough 1976; Gosling 1982, Guarneros et al. 2020). For example, horses spontaneously exploring or ridden in a familiar arena nevertheless make a record of objects and feces located on the surface of the arena by sniffing and they show that they remember objects on that day by not returning to sniff them again (Burke and Whishaw, 2020; Whishaw and Burke, 2021). This ability to recall objects in relation to both space and time is indicative of episodic-like memory; that is, knowing “where, when, and what” (Clayton and Dixon, 1988; Clayton et al. 2001, Marshall et al. 2013, Tulving 2016). That this form of episodic-like memory may be spontaneously displayed by horses raises the question of what is remembered and what is the memory duration of remembered items. In a previous study of memory duration for dung deposits, object location was recorded but sniffing duration, which might indicate memory strength and interest in items, was not measured (Burke and Whishaw, 2020; Whishaw and Burke, 2021). Therefore, for the present study, the testing procedure was modified so that location and sniff duration could be measured concurrently.

Rather than examine the behavior of horses at liberty or when ridden, the experimenters obtained experimental control by leading the horses from a starting location to a point at which they could see a target object that they could then spontaneously approach and investigate. This procedure allowed the presentation, location and identity of the target to be systematically varied. Trials were video recorded so that sniffing behavior and its duration could be subsequently assess by frame-by-frame analysis. For the study, many of the same horses were tested in their home indoor or outdoor arenas in four different locations over a two-year duration.

## Methods

### Subjects

Nineteen domestic horses (9 geldings, 2 stallions, 8 mares), aged between 4 and 21 years, were used for the study. Every horse was not available for all tests and so the number of horses used varied and was smaller than the total count used in each experiment. Testing occurred at five acreages: 3 in Lethbridge, Alberta (n=13 horses), 1 in Fernie, British Columbia (n=3 horses), and 1 in Lyn, Ontario (n=2 horses). The horses were tested in indoor and outdoor riding arenas with which they were familiar between the months of August 2020 and August 2022. There were no obvious difference in the pattern of behavior displayed in the indoor and outdoor arenas or in different arenas and so test location was not a dependent variable. Most of the horses participated in a number of tests, with test separated by at least 30 days or longer.

### Video recording

The object inspection behavior of the horses was recorded with an iPhone 11 and the video records were downloaded onto a MacBook Pro and/or Samsung Galaxy for analyses. For some experiments, a photographer sat in front of the dung stimulus, at a distance of 3 m, to obtain a face on view of behavior. For the experiment in which blinking was recorded, the iPhone was mounted on a stand located 1 m to the side of the stimulus and was remotely activated. The positioning of the photographer or camera did not obviously affect behavior. Video inspection for Frame-by frame-analyses were made using Quick Time Player (Apple Computer) and DJV2 (GitHub) programs.

### Stimulus Plate

The stimulus samples were present to the horses on 30 cm diameter silver colored plastic plates. Prior to each test, the horses were presented with a clean plate by holding it up so that they could sniff it. Identical plates were used for tests that required a number of stimulus samples. The dung samples were heaped on a plate so that only the outer rim of the plate remained exposed (see photos in Figure 1).

**Figure 1.**
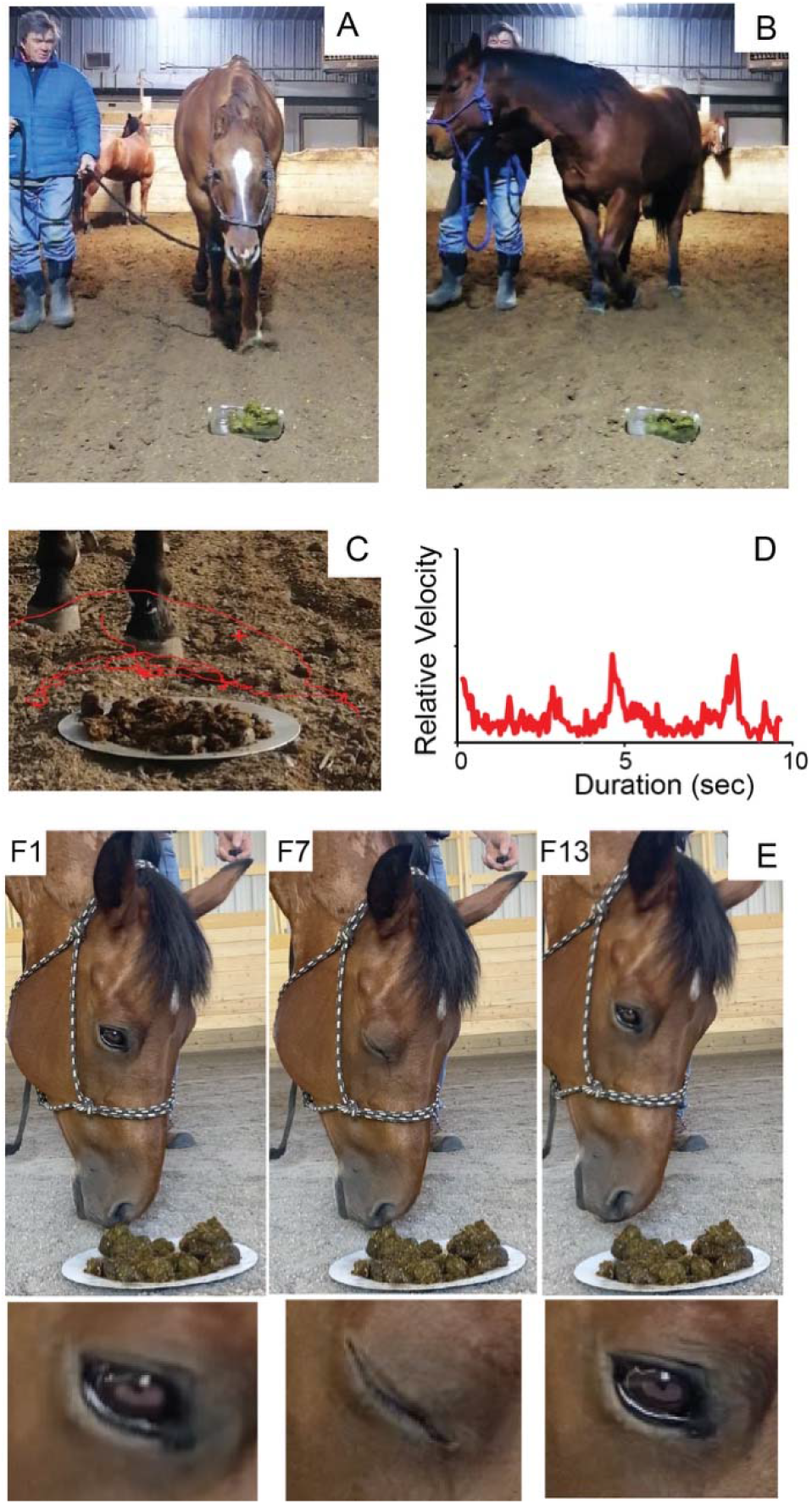
A. Approach. Horses were led on loose lead from a tie up spot on the wall of the arena to the proximity of the dung stimulus and were then allowed to walk forward on their own. B. Blocking. A horse that had previously sniffed the dung refuses to approach by stepping in front of the handler. C. A kinematic trace of the center of a horse’s nose over the dung during a sniffing episode. D. The velocity of nose movement of the trace shown in C, in which periods of sniffing are interspersed with larger head movements, signified by high velocity peaks, across the dung. E. A disengage-blink by a horse as it lifts its head from sniffing (Frames F1, F7, F13 from nose removal onset) with close-ups illustrating a complete eye blink.

### Stimuli

Two types of stimuli were used, dung samples and mud samples. The mud samples were arranged on the plates so that they had a similar visual appearance to the dung samples. Dung samples were either fresh or days old, depending upon the experiment. Some dung samples that were reused were enclosed in a zip-top bag and the bag placed into a 1.7L food-grade glass container with an airtight snap-and-lock lid.

In a preliminary study with 4 horses, dyads of dung stimuli were concurrently presented side by side and separated by 20 cm for a comparative assessment. Dyads consisting of a horse’s own dung, the dung of a congener with whom the horse was stabled, or dung of an unknown mare or gelding. The subjects were allowed to investigate the samples twice with a 5-minute wait between each trial. Comparisons of sniff duration showed that a significantly (p<0.05) higher proportions of total trial time was spent sniffing unfamiliar stimuli (gelding or mare) than familiar stimuli (own or pasture-mate). These findings are in accordance with Krueger and Flauger (2011), who find that horses distinguish broadly between familiar and unfamiliar conspecifics. Therefor for subsequent experiments, the source of the dung is identified.

### Arena Setup

Prior to testing, all objects were removed from the arena and horses were familiarised with the area by being tied up for at least 20 minutes. The testing plates were placed on the ground at least three metres from a wall. For single day tests, a fresh dung sample was placed on the plate and for some repeated tests the same dung was used. A handler led each horse towards a plate on a loose rein and once the horse noticed the plate, the tester stopped allowing the horse to walk forward on its own to investigate the stimulus.

## Behavioral measures

### 1. Approach

An approach was counted if a horse freely walked forward and brought its nose in proximity to a stimulus as if to sniff it. An approach was not counted if a horse stopped did not advance within 10 sec, veered to either side, or walked over the stimulus without lowering its head.

### 2. Sniff duration

Sniff duration was measured by counting video frames (30 f/sec) and converting the counts to seconds. Frame counts began when the nose of a horse stopped its descent toward a stimulus and ended with the first frame indicating the horse was lifting its nose away from the stimulus.

### 3. Sniff laterality

A measure of laterality of sniffing was made by counting of frames in which either the left or right nostril was clearly preferentially directed toward a dung stimulus or was moving across the dung stimulus. The measures were made from 15 horses (5 mares, 10 geldings) given two tests. The first test consisted of a novel dung sample and the second test was with the same sample after a delay of 20 min. Results for the left and right nostril were summed and subjected to a t-test for correlated groups.

### 4.. Ear laterality

Ear position was measured by counting frames on which either the left ear or the right ear was relatively more forward. The measures were made from 15 horses (5 mares, 10 geldings) given two tests. The first test consisted of novel dung sample and the second test was with the same sample after a delay of 20 min. Results for the left and right ear were summed and subjected to a t-test for correlated groups.

### 5. Blink disengage

A blink-disengage was counted if a horse briefly completely closed its eyes as it stopped sniffing the stimulus and raised its head away from the stimulus. Counts of blink-disengage were made from 12 horses (8 geldings and 4 mares) sniffing either mud samples or dung samples at three different intervals, giving a total of 72 trials. The measure of blink-disengage was the total incidence recorded from each horse.

## Procedure

### 1. Comparison of dung and mud

Thirteen horses (9 geldings and 4 mares) were tied up in pairs in an outdoor arena for 20-min to adapt them to the arena on each of three testing sessions. In each session, the horses were presented with a plate at one location and then 20-min later with plate at a second location. Locations were separated by 7-m from each other. One plate contained dung from an unfamiliar gelding and the other plate contained an equal sized similarly appearing pile of moist mud. The test procedure was repeated at intervals of 0, 6 and 24 hours. The stimuli were placed in a zip-top bag and the bag was placed into a 1.7L food-grade glass container with an airtight snap-and-lock lid between tests.. Seven of the horses were taken to the dung stimulus first and 20-min later they were taken to the mud stimulus, for the other 6 horses the order of presentation was reversed.

### 2. Comparison of similar and different dung at different locations

Fourteen horses (6 mares and 8 geldings) were tied up in pairs in an arena for 20-min to adapt them to the arena. They were then given 4 trials, with 20-min intertrial intervals, in which they were taken twice in succession to one location and twice in succession to a second location. The two locations were separated by 7 m. In a first test, the horses had examined the dung at one location, the dung was moved to the second location, so that a horse reencountered the same stimulus but at a different location. In a second test, repeated about 30 days later, the dung at the two locations was different.

### 3. Comparison of 20-min and 24-hr intervals

Four horses, three adult geldings and one mare that lived together as a herd, were given two tests in which they were presented three different dung samples in succession. The stimuli for Test 1 were arranged in a triangle pattern, 20-m apart, at the south end of the arena and the stimuli for the Test 2 were arranged in a triangle pattern, 20-m apart, at the north end of the arena. For each test, the presentations were given 6 times in succession. The interval between the six successive presentations of the 3 stimuli was 20-min for Test 1 and 24-hr for Test 2. The 24-hr test was given 30 days before Test 2. At the completion of the tests, one further stimulus presentation was given with the intervals reversed, so that after the last trial of Test 1, the horses were tested once more 24-hr later, and after the last trial of Test 2, the horses were tested again 20-min later. For both tests, the 3 dung samples were fresh for the first trial, but the samples and their location were not changed during the remainder of the trials of that test. The sources of the dung samples placed at each location for the tests were different. *Location 1*, the dung sample was from an unknown gelding; *Location 2*, the dung sample was from one of the geldings in the herd; *Location 3*, the dung sample was from an unknown mare. For Test 1 (24-hr test), the horses were haltered before each trial and for Test 2 (20-min test) horses they were haltered and tied up in the test area between trials. The order in which the horses visited the three location was 1-3 for two horses and 3-1 for two horses.

### 5. Twenty-four-hour retention test

Nine horses (6 geldings and 3 mares) were given two tests to assess their retention of a dung sample obtained from an unknown gelding. In Test 1 they were taken to one dung sample repeatedly at 5-min intervals until they no longer approached it or sniffed it. Then, 24-hr later the procedure with the same sample was repeated. In the interval, the dung sample was placed in a zip-top bag and the bag was placed into a 1.7L food-grade glass container with an airtight snap-and-lock lid. The procedure for Test 2 was similar except that a fresh dung sample from the same horse was used for the 24-hr test. The two tests given with the starting point and the dung location in the same place, but the tests were separated by about a 30 day interval.

## Results

### 1. General observations

#### Stimulus approach and inspection

When the horses were led to the staring position near a stimulus, they always detected the stimulus at a distance greater than about 3-m and then spontaneously approached the stimulus with head lowered and ears forward. Thus, on the approach and during sniffing, the handler was standing back at about the midpoint of the horse’s body (Figure 1A). On only three occasions, in all experiments, did the handler have to encourage the horse to walk forward to the stimulus within a distance closer than 3 m. If the horse had been already taken to a given dung sample on the same day, it might fail to approach the stimuli by walking passed it, turning away, or even turning to the side to block the handler (Figure 1B). Two horses, a gelding and a mare, were notable for displaying blocking, but various degrees of turning into or away from the handler were displayed by many horses on retention tests. (Blocking behavior is a recognizable behavior in horses, as a mare might block a foal from approaching another horse.)

#### Sniff behavior

Upon reaching a dung stimulus, a horse would lower its head so that its nostrils were just above the stimulus. The horse then made sniffing movements as it moved its nose back and forth across the stimulus (Figure 1C). Sniffing occurred in sniff bouts, restricted to a small region of the stimulus, interspersed larger head movements, in which the nostrils were moved to a new location on the stimulus (Figure 1D). Sniffing of dung samples that were fresh were associated with only light brief nose contacts on the dung. Sniffing dry dung or mud stimuli were sometimes associated with some horses nosing the stimuli or nibbling it (*coprophagia*). No horse was observed to engage in Flehmen, an olfactory-related behavior in which the animal raises its nose and curls its upper lip to expose the upper incisors after sniffing (Eisenberg and Kleiman, 1972; Jezierski et al, 2018).

#### Blink disengage

When a horse disengaged a stimulus by lifting the nose or turning it away to one side, the disengage movement was usually associated with a blink (Figure 1F). Of 12 horses that were video recorded from the side on a total of 72 trials, the eye was visible on the video record on 61 trials (the head was turned away or the mane covered the eye on the remaining trials). Of these 61 trials, a blink was observed on 58 trials. Of these blinks, 53 were associated with complete eye closure and 5 were associated with partial eye closure. The first video frame on which a disengage movement on the nose was observed was designated as the “0” frame, and the first movement of the blink was identified in relation to this point. The average frame at which the blink began was frame − 0.275±0.6, indicating the disengage of the nose and the onset of the blink were almost concurrent.

#### Sniff laterality

An assessment of nostril preference for sniffing was made for 15 horses that sniffed a dung stimulus on a first presentation and second presentation 20 min later, with the observations collapsed across tests. The t-test for correlated groups that assessed the duration for the left vs right nostril preference in relation to proximity to the dung gave no difference in nostril duration directed to the dung, t (14)=0.71, p>0.05.

#### Ear laterality

An assessment of nostril preference for sniffing was made for 15 horses that sniffed a dung stimulus on a first presentation and second presentation 20 min later, with the observations collapsed across tests. As assessed by the ear that was preferentially forward during dung sniffing, there was no preference in ear position, t(14)=1.25,p>0.05

## Experimental results

### 1. Horses sniff dung longer than mud

Figure 2A diagrammatically shows the location of the dung and mud stimuli and Figure 2B illustrates the mean sniff duration made by the horses to the two stimuli. The horses approached and sniffed both the mud and dung stimuli, and an ANOVA indicated that the horses sniffed the dung for a longer than they sniffed the mud, Stimuli F(1,12)=15.5, p=.002, *η^2^p* =0.564. A comparison of sniff durations at the three different times, 0, 6 hr and 24 hr, gave a difference, Test time F(2,24)=7.6, p=.003, *η^2^p*= 0.389. Follow-up Bonferroni tests indicated that sniff duration was longer to the dung on the immediate test vs the 6 hr test (p<0.001) and longer on the 24 hr test than on the 6 hr test (p<0.05). The analysis also gave a significant Test time by Stimulus effect, F(2,24)=6.13, p=.007, effect. This was accounted for by the relatively unchanging sniff durations as a function of test interval to the mud vs the changing sniff durations as a function of test time to the dung. There was also no significant sex difference, F(1,11)=1.08, p=0.31.

**Figure 2.**
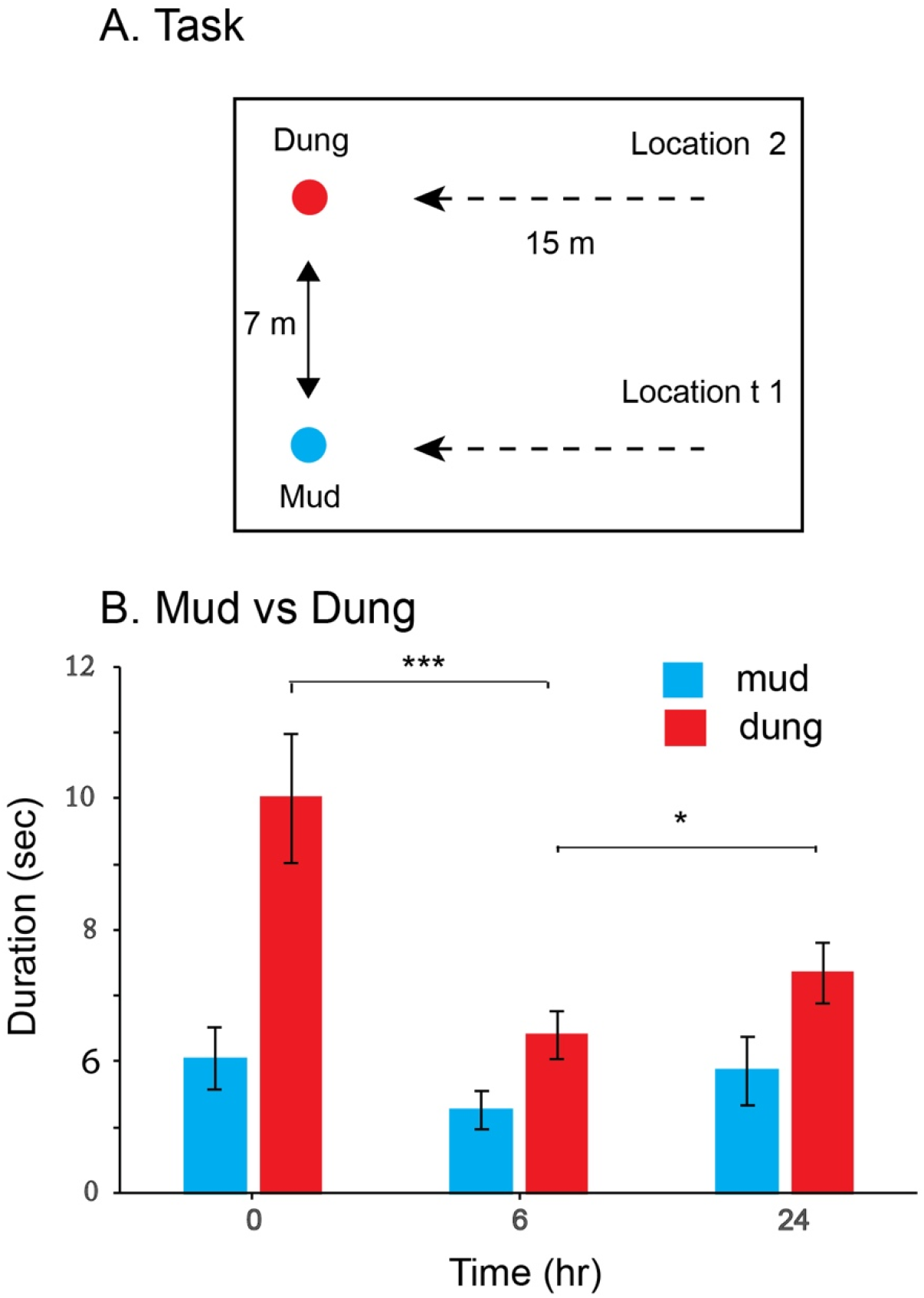
A. Task used to compare dung and mud sniffing. The start point was 15-m away from the stimulus and the stimuli were separated by 7-m. Half of the horses approached the dung stimulus first and half approached the mud stimulus first. B. Sniffing durations (mean and se) showed horses sniffed dung significantly longer than mud, showed decreased sniffing of dung a 6-hr and an increase at 24-h. ***=<0.001; *=<0.05

### 2. Horses remember dung upon rencounters at new location

The experiment examined whether horses remember a dung sample when they reencounter it at a new location. Each horse first approach dung at Location1 for two successive trials and then approached dung at Location 2 for two successive trials. For Test 1, the same dung was at Location 1 and 2 and for Test 2, different dung was at Location1 and Location 2 (Figure 3A). Were the horses to remember the dung independently of its location, a decrease in sniff duration would be expected when they reencountered familiar dung at a new location. This prediction was confirmed as shown in Figure 3B and supported by the ANOVA comparing sniff duration as a function of test, trial and location. When horses reencountered the same dung at a new location their sniff duration decreased relative to the duration of sniffing when encountering new dung at a new location. Thus, there was significant effect of Test, F(1,12)=9.52, p<.01, η^2^p=0.4; Location (F(1,12)=5.68, p=0.35, *η^2^p*=.32; and Trial, F(1,12)=64.8, p=0.001, *η^2^p*=.84 and Test by Trial by Location, F(1,12)=5.14, p=).43, *η^2^p*=0.30. The pattern results was the same for mares and geldings, but the analysis also indicated that the sniff durations for the mares was shorter than for the geldings, Sex F(1,12)=11.03, p=0.000, *η^2^p*=0.05; Figure 3 C).

**Figure 3.**
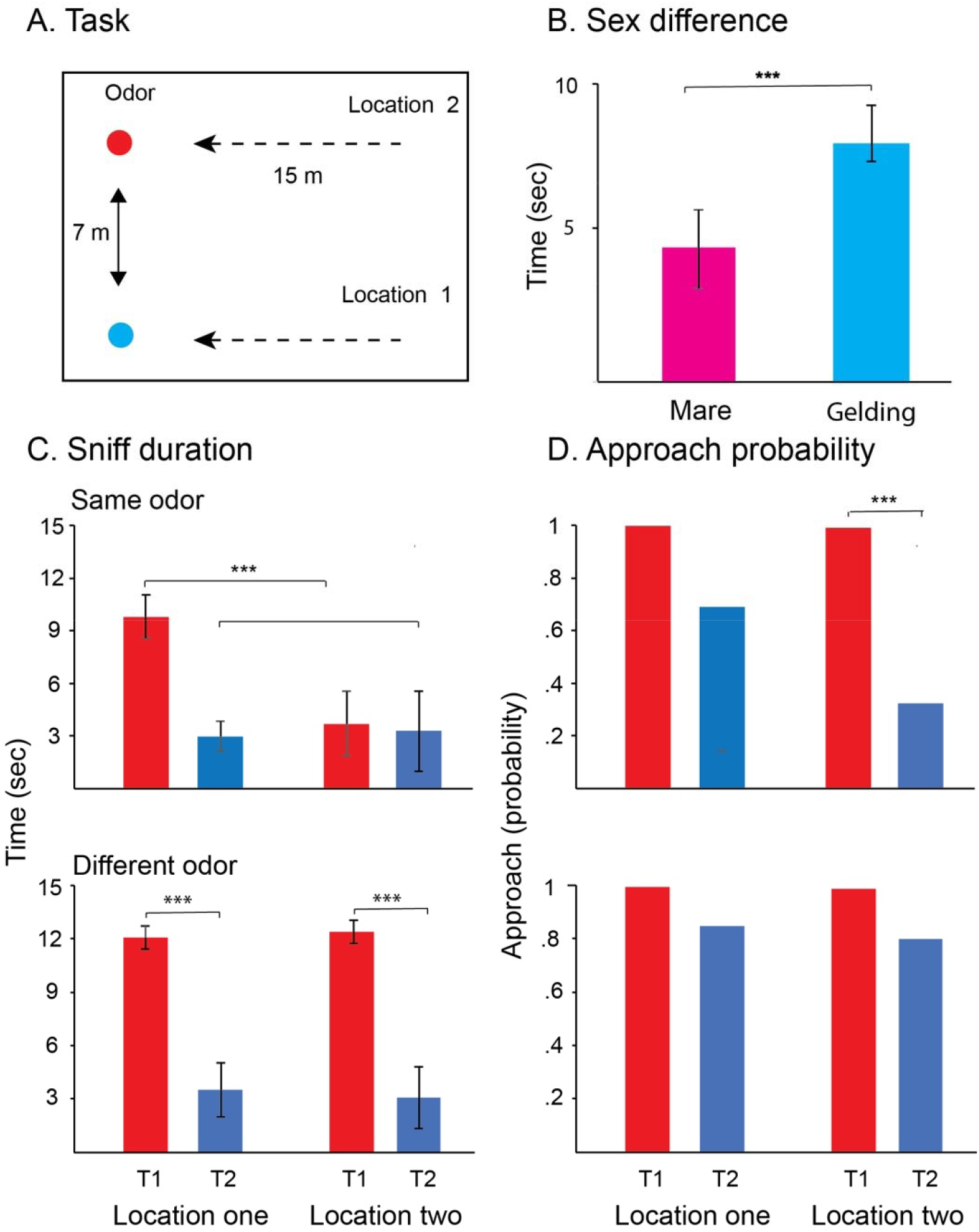
A. Task used to compare same-dung and different-dung sniffing. The start point was 15-m away from the stimulus and the stimuli were separated by 7-m. For one test, the sample was at Location 1 and horse encountered the same dung at Location 2. For the second test, the dung at Location 2 was different. B. Across all measure mares sniffed for shorter durations than geldings. C. Sniff duration in same and different dung tasks gave a significant interaction showing that horses remember the sample in the same dung task. D. Approach probability measures gave a significant interaction showing the approach probability decrease in proportion to previous sample exposure irrespective of location. ***=<0.001

The measure of approaches gave similar results to that obtained with sniff durations (Figure 3D). Overall, the horses approached the dung less frequently going to the same location on Trial 2 than they did on Trial 1, Approach F(1,12)=38.94, p<0.001, η^2^p=0.78, indicating that they remembered their experience of sniffing dung at that location on the previous trial and chose not to sniff it again. There was a significant effects of Location F(1,12)=12.93, p<0.01, η^2^p=.54, and Test (1,12)=8.09, p<.05, η^2^p=.42, as might be expected, as the horses sometimes did not approach dung a second time at the same location but always approached dung at a new location, Test by Trial F(1,12)=5.51, p<0.05.

### 3. Horses remember dung withing a day but not between days

To determine whether horses remember dung samples at specific locations over days, horses were taken to three successive dung samples at either 20-min intervals or 24-hr intervals. A map of the test arrangement shown in Figure 4A, illustrates the path taken for the 24-hr test interval given at the south (S) end of the arena and the 20-min test interval given at the north (N) end of the arena. A summary of the results comparing a 20-min retest interval to the 24-hr retest interval is given in Figure 3B. There was a significant test interval effect, with shorter sniffing durations obtained on the 20-min test compared to the 24-hr test, Duration F(1,3)=33.87, p=0.01, *η^2^ p*=0.919. There was also a significant effect of test day, with a decline in sniff durations occurring over trials for the 20-min test interval with no decline in duration over trials for the 24-hr test interval, Days F(1,3)=209.7, p<0.001, *η^2^ p*=0.986; Test by Days F(1,3)=3.95, p=0.14, *η^2^ p*=0.569. As is shown in Figure 4B, when the test intervals were reversed after the 6 trial, there was a significant decline in sniff durations for the interval change from 24-hr to 20-min and a significant increase in sniff duration of the interval change from 20-min to 24-hr, Tests F(1,3)= 42.37, P<0.007, *η^2^p*=0.934; Time F(1,3)=5.66, p=0.98; Test by Time F(1,3)=18.21, p=0.24, *η^2^p*=0.859.

**Figure 4.**
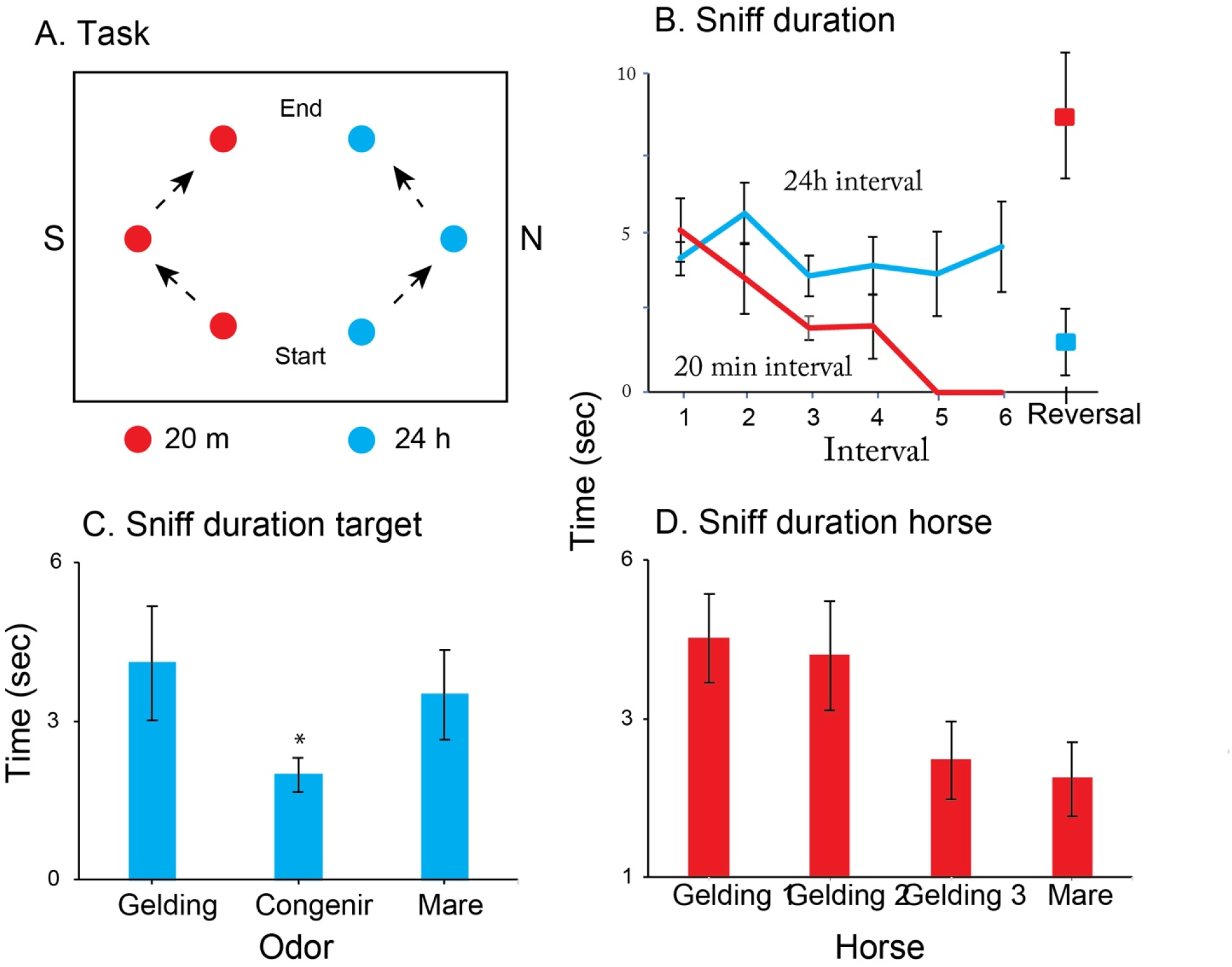
A. Task. The horses were taken 6 times to 3 dung samples in the south end of the arena at 24-hr intervals and 6 times to 3 dung samples in the south end of the arena at 20-min intervals. Two horses visited samples in the direction of the arrows and for two horses the direction was reversed. B. Sniff durations remained high with trials at 24-hr intervals and declined with trials at 20-min intervals. Durations reversed when the time interval for the groups was reversed. C. Across both tasks, the sample that was taken from one of the herd geldings was sniffed less than the samples taken from an unknown mare or gelding. D. Sniff durations by individual horses were distinctive.

Because different samples were used at each of the locations, sniff durations at the different locations were compared and are shown in Figure 4C. An analysis of samples did not give an effect of Location, F(2,6)=3.81, p=0.086, *η^2^ p*=0.482, but the Day by Location effect was significant, F, F(10,30)=2.63, p=0.02, *η^2^p*=0.882, mainly due to the shorter sniff durations at the conspecific dung sample at Location 2. The between subjects analysis was also significant, F(1,3)=22.407, p=0.18, *η^2^ p*=0.882, with two geldings displaying consistent and equally long durations and the third gelding and the mare displaying consistent and equally short durations (Fig. 4D).

### 4. Repeated dung stimulus encounters do not result in long term retention

When the horses were allowed to walk repeatedly to a dung sample (Figure 5A), they displayed prolonged sniffing on the first trial but thereafter sniffing duration decreased in a trial-by-trial pattern until within a few trials most horses no longer approached the dung. On a retention test given 24-hr later, there was seemingly no recollection of the previous test. On the 24-hr test, the horses again inspected the dung for a similarly long period on their first trial and again displayed decreased trial-by-trial sniff duration until they no longer approached the dung. This pattern of behavior occurred on the test in which the sample of dung was used for both the initial test and the retention test and for the test in which a fresh dung sample from the same donor was used for the second test (Figure 5B left vs Figure 5B right). The ANOVA confirmed these results by giving a significant effect of Trial, F(1,7)=30.9, p<0.001, *η^2^ p*=0.82, but no significant effect of Test, F(1,7)=3.26, p=0.14, *η^2^p*=0.32 or Sample by Trial Interaction, F(3,21)=0.21, p=).89, *η^2^p*=0.03. The sniff durations made by individual horses on the two tests were significantly correlated, r=0.79, F(1,8)=11.6, p=0.01, see Figure 5C). Although the mares had overall shorter sniffing durations that the geldings, the group differences did not reach significance, F(1,7)=2.57, p=0.15, *η^2^p*=0.27.

**Figure 5.**
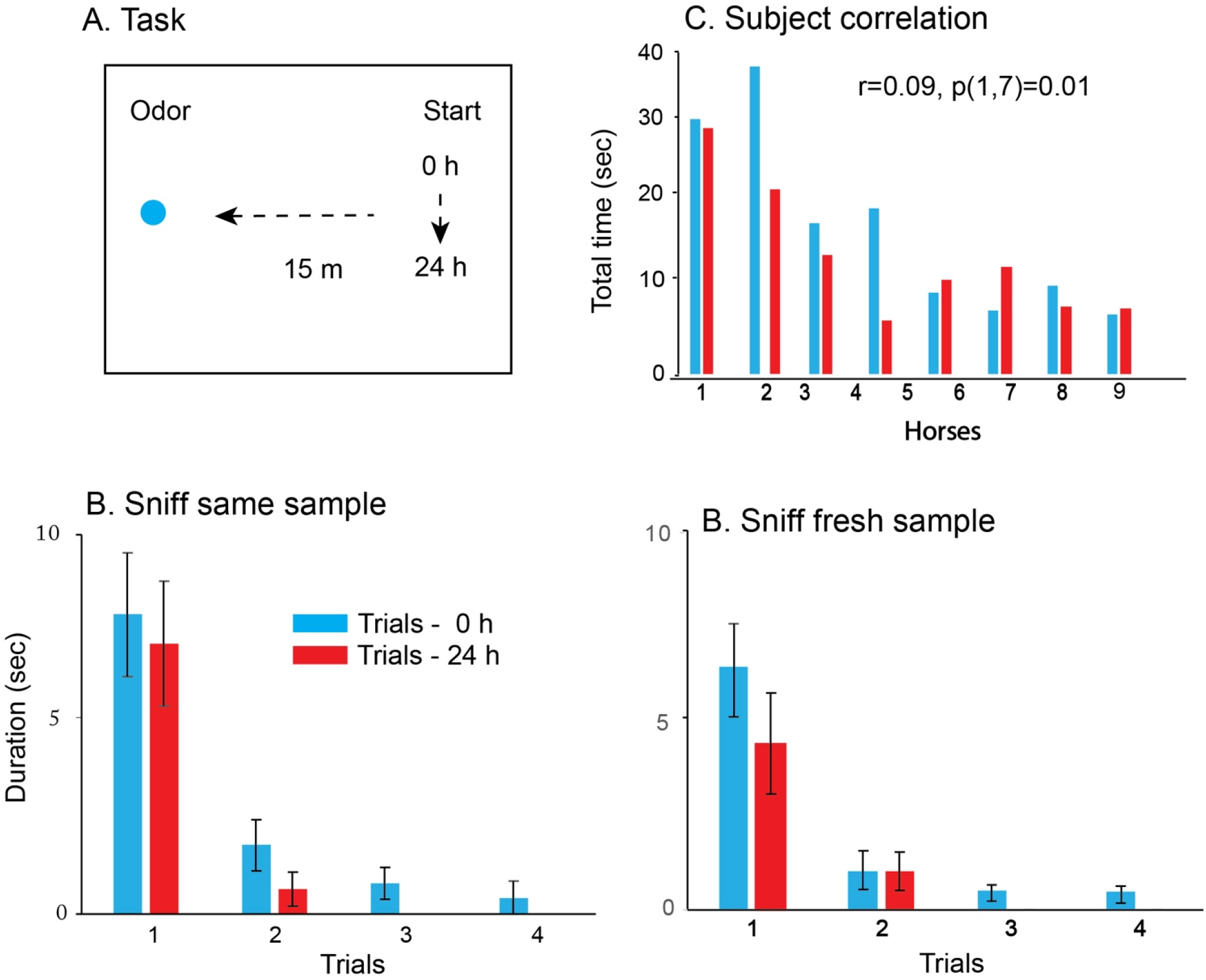
A. Task. Horses were taken to one dung location repeatedly until they no longer approached, and 24-hr later given the same test with the same sample or with a fresh sample from the same horse. B. Sniff durations for the initial sample and on the retention at 24-hr were the same for the same sample (left) or a fresh sample from the same horse (right). C. Total sniff durations by the horses in the two tests were correlated. Note: long sniff durations on the first trial of each day.

## Discussion

This study assessed horse memory for dung samples by measuring approaches to dung piles and sniffing duration of the dung as a function of exposure number and time interval. Horses sniffed dung for longer times than non-dung items and sniffed dung from unknown horses for longer times than from known horses. Intraday approach and sniffing duration decreased rapidly with repeated exposure to a sample, supporting the conclusion that horses display excellent episodic-like memory for the location and identity of dung deposits on a given day. By contrast, the results of four experiments, gave no similar interday retention. This pattern of results suggest that horses use a day-to-day time-limited episodic-like memory of dung location and identity as part of their representation of locations that they occupy. Thus, memory duration can be influenced by ecological constraints.

In previous studies of dung sniffing behavior, horses were tested either at liberty (Krueger and Flauger, 2011) or under saddle (Whishaw and Burke, 2021), whereas in the present study they were lightly controlled on a lead line. This procedure was accepted by all horses as none passed up the opportunity to sniff dung samples. As performance horses, all had experience with handling on a lead line, giving the handler good control of the horses’ movement (Hill, 2012). The horses were, however, more likely to return to an object as second time than was observed when they were free to move about on their own (Whishaw and Burke, 2021). This tendency to return to a stimulus when taken proximal to it on a lead line proved beneficial because for most experiments relative approach and sniff duration provided a measure of spatial and object memory. Nevertheless, the horses’ approach to samples were not forced, because after a handler brought them within visual proximity, they moved forward on their own, allowing the handler to stand behind them as they inspected the object. In it noteworthy that the procedure is not different than used with tracking dogs (Johnson, 2003), and could be readily used for other horse object recognition studies.

Difficulty in defining episodic memory in nonhuman animals has led to the heuristic of using the term episodic-like for the nonhuman forms of episodic memory, memory representing the what, where, and when of items (Clayton and Dixon, 1988; Clayton et al, 2001; Olsson and Brown., 2006). Results from previous studies with horses, along with the results of the present study, are consisted with the idea that horse memory for dung encounters has features that allow it to be considered a form of horse episodic-like memory (Whishaw and Burke, 2021). The memory of dung items is formed without a definite decision, as horses are interested in sniffing dung to the extent that they are unlikely to pass up an opportunity do. Horses remember where they have sniffed an item because when encountering an item that they have previously investigated at a location, they may pass up the opportunity to re-examine it and if they sniff it again, their sniffing duration is reduced. Horses also remember the identity of the item that they have sniffed, even if they find it at a new location, because as shown in the present study, upon reencounters, their sniffing duration is shorter. The intraday memory duration of individual dung encounters does not appear time dependent. Memory for dung was retained equally after minutes or hours, and in the present study, it was as good at 20-min as it was at 6-hr. This duration is certainly long enough to be described as episodic-like memory (Schwartz and Evans, 2001). Taken together, this evidence suggests that horses display a form of episodic-like memory in relation to dung that has some of the features of episodic-like memory described for olfactory odors by dogs and rats (Lo et al, 2020; Quaranta et al, 2020). It also has features of episodic-like memory described for food items in other animal species, including honeybees, scub jays, pigeons, hummingbirds, rats, and primates (Clayton et al, 2001)

Nevertheless, unlike episodic-like memory described for other species in other tasks, the memory for dung items by horses does appear to have a temporal index. When returning to a dung item after a day’s delay, the retained duration of a horse’s sniffing suggests little to no memory of a previous visit. In the four experiments described here, horses taken daily to the same dung location displayed increased sniff durations relative to intraday dung reencounters. Even when horses were taken repeatedly to a dung item until they refused to approach it on intraday tests, they showed prolonged sniffing on the first encounter on a subsequent interday test. This finding was obtained when the dung item was preserved by wrapping and, to control for the possibility that there were qualitative changes in the preserved dung, when fresh dung from the same donor was used. The interday recovery of dung sniffing confirms a previous study that found that horses under saddle seldom returned to objects that they had sniffed on that day but did return to sniff them when taken to the same arena on a subsequent day (Whishaw and Burke, 2021).

An explanation for the seeming interday forgetting of dung items may relate the ecology of horse movement. Horses in open ranges move long distances as they forage and in doing so are likely encounter other horses (McCourt, 1984; Ransom and Cade, 2009; Ranson and Kaczensky, 2016). Thus, it is likely that horses are motivated assess/reassess each area as they move into it in relation to potential conspecific occupancy or other threat. Dung sniffing is a useful component of occupancy area inspection, as it provides information about the recent presence of other horses (Feist and McCullough, 1976; McDonnell, 2000; Saslow, 2002). Information contained in dung could include threat potential, food consumption of the donor, sexual status and so forth (Galef et al, 1994; King and Gurnell, 2006; Krueger and Flauger, 2011; Péron et al., 2014, Whishaw and Burke, 2021). For these reasons it would be important for horses to assess dung in area that they enter. Other evidence of interday forgetting by horses that may also be indicative of risk-assessment comes from the shying behavior of horses. It is recognized by horse riders that horses will persistently shy at objects to which they have been seemingly habituated to the previous day (Budiansky, 1997). Thus, their daily renewed curiosity about dung and wariness about objects may both be related to staying safe as they move about.

Study of horse laterality suggested that horses display a nostril preference for sniffing (Larose et al, 2006; McGreevy and Rogers, 2005; Siniscalchi, 2015). Such a preference did not occur with dung sniffing as measured here. The horses usually scanned a dung item extensively, such that one then the other nostril was more proximal to the dung. It is likely that dung is rich in olfactory information and it could not be determined from the present analysis whether different information is being extracted by each nostril as sniffing progresses. Similarly, there was no consistent ear orientation displayed during sniffing, except that when the horses approached a dung item their ears were forward and as they began to sniff their ears usually became relaxed. The horses did display two kinds of blinks, partial and complete, during dung sniffing and they nearly always displayed a complete blink as they disengaged from sniffing. Disengage-blinks have been described in anthropoid primates including humans after period of visual attention (de Bruin *et al*. 2008; Sacrey *et al*. 2011; Hirsche *et al*. 2022), but this may be the first description of a blink after a period of olfactory attention. It is interesting that fMRI evidence has suggested that blinking may be a sign of brain network change, as might occur when going from an alert to resting state or shifting attention from one item to another (Nakano et al, 2013). A disengage-blink by the horse may suggest a change in brain sensory networks used for olfactory attention to a network dependent on vision related to moving away from the dung sample.

In other work, it has been noted that there are sex differences in investigating stud piles as well as individual differences in that stallions display more investigatory behavior than geldings (Feist and McCullough, 1976; McDonnell, 2000; Saslow, 2002). In the present study, equally large samples of mares, geldings and stallions were not available, but it was noted that the sniff durations of the mares were lower than that of the geldings, although because of limited samples statistical differences could not be adequately assessed in all experiments. It was also noted that there were large individual differences between geldings, and future work could consider the source of these differences.

In conclusion, human episodic memory is not defined by duration as most are forgotten as soon as they are acquired, but they can endure and they are discussed with respect to utility for mental time travel (Baudry, 2020; Ruby and Clayton, 2012). The episodic-like memory for dung items described here for the horse may serve a function in maintain social cohesion and safety. Horses form relatively small herds that are stable but move around a good deal where they may encounter other horse herds. Dung sniffing may be a source of information about who is around and who might pose a threat. Stallions have the most interest in maintaining herd cohesion and they are reported to be most interested in dung sniffing, and here we find that geldings are more interested in sniffing than mares, although all horses sniff dung. The temporal boundaries for memory about dung, in which items are remembered for a day and reinspected after a day, likely reflect the ecology of herd movement. Information provided by dung is most relevant as horses move through feeding and resting areas, which they most likely do daily. This interpretation of the utility of the episodic-like memory of dung sniffing in the horse suggests that if it does have any utility for mental time travel in the horse, it may be limited to the relatively brief duration of a day.

